# U1 snRNP suppresses microRNA biogenesis by alternative intronic polyadenylation in melanoma

**DOI:** 10.1101/2022.06.01.479622

**Authors:** Sandra Vorlova, Gina Blahetek, Ruggero Barbieri, Yvonne Kerstan, Annabelle Rosa, Maja Bundalo, Manuel Egg, Elke Butt, Erik Henke, Lars Dölken, Roland Houben, Bastian Schilling, Utz Fischer, Florian Erhard, Alma Zernecke

## Abstract

Activation of intronic polyadenylation signals results in premature cleavage and polyadenylation (PCPA). The majority of mammalian miRNAs are also located within intronic regions of protein-coding genes and are transcriptionally co-expressed with their host genes. Here we show that U1-dependent PCPA by telescripting dysregulates miRNA biogenesis. When U1 is reduced, miR-211 levels are decreased as a direct consequence of activation of a newly identified alternative intronic polyadenylation signal located upstream of miR-211 within its host gene TRPM1. Various melanoma cell lines revealed decreased U1 levels and a shift from full-length to truncated TRPM1 isoforms with concomitant decreased miR-211 expression. Modulation of TRPM1 alternative polyadenylation (APA) by morpholino oligonucleotides inhibits and potentially restores miR-211 expression to endogenous levels. This mechanism of intronic PCPA and its effects on miRNA biogenesis represents a previously unrecognized layer of gene expression regulation suitable for therapeutic modulation.

**Graphical Abstract:** 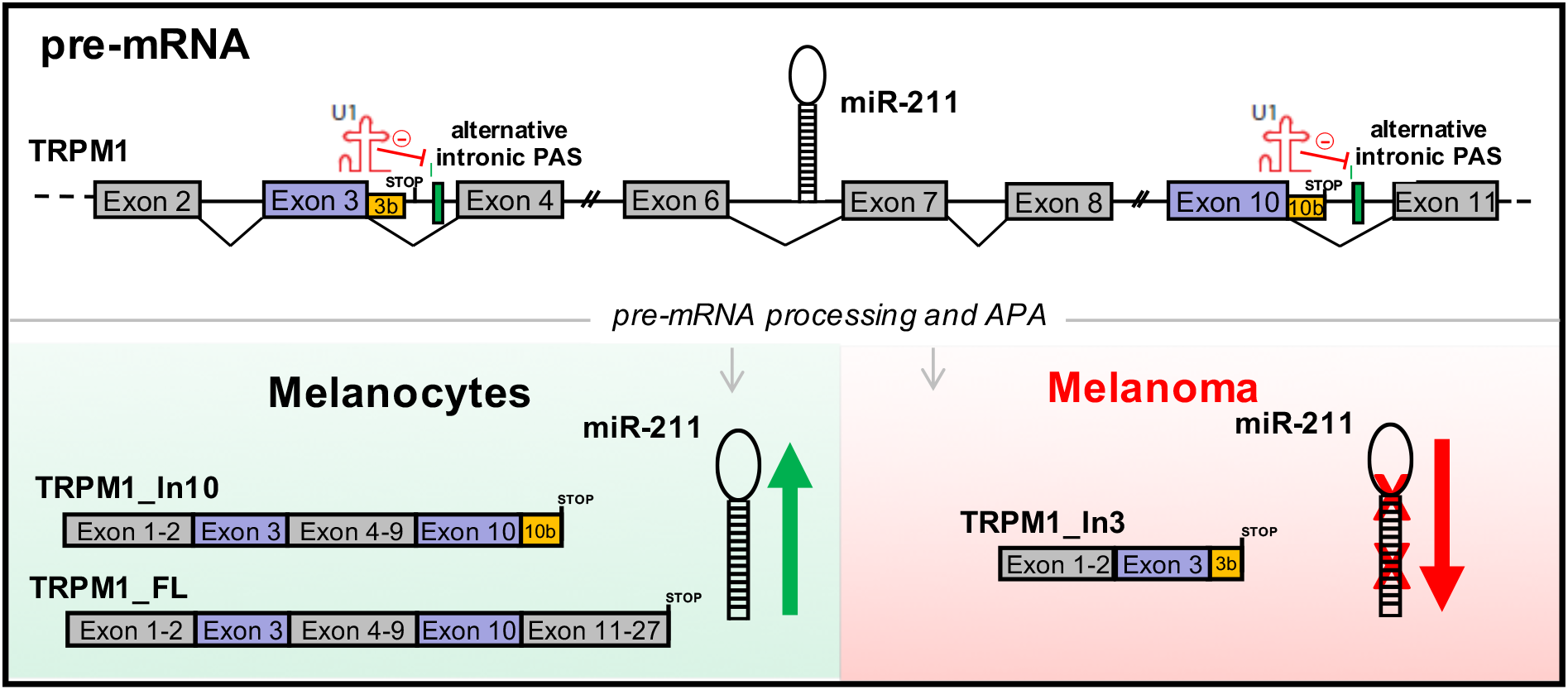

## Main

Messenger RNAs (mRNAs) undergo important post-transcriptional modifications that have profound effects on the resulting transcripts. It is estimated that ∼70-75% of human genes contain alternative polyadenylation signals (PAS) ^1^, and alternative polyadenylation (APA), i.e. activation of these alternative PAS, is emerging as a major player in controlling gene expression ^2,3^. Recent studies have shown that poly(A) site usage is frequently altered in immunological, haematological and neurological diseases, as well as in cancer ^4-7^. Genome-wide PCPA due to activation of APA has also been described as the predominant and rapid heat shock response ^8^. Usage of alternative PAS located within the 3’untranslated region of a gene results in transcripts with reduced length of their 3’UTR, elimination of microRNA-bindings sites, and consequently increased expression ^9^. On the other hand, activation of PAS located in internal introns results in stable and shortened mRNAs by PCPA and the generation of C-terminal truncated protein isoforms with often antagonistic functions ^10^. U1, which is an essential part of the spliceosome and the most abundant non-coding small nuclear RNP in vertebrates, inhibits activation of cryptic or alternative PAS ^10,11^. Furthermore, U1 suppresses malignant activity as its inhibition leads to an invasive and migratory phenotype of cancer cells ^12^. Intriguingly, the majority of mammalian microRNAs (miRNAs) are also located within intronic regions of protein-coding genes ^13,14^, and miRNA expression largely coincides with the transcription of their host genes, indicating co-regulation and generation from a common precursor transcript ^15^. Dysregulation of miRNA expression is particularly closely associated with human cancer ^16,17^, substantiating the need for miRNA based therapeutics to modulate miRNA activity ^18^. Melanoma, the deadliest type of skin cancer, develops from epidermal melanocytes ^19^. Despite advances in therapeutic options for metastatic disease in recent years ^20^, resistance to therapy remains a major clinical challenge in melanoma. New insights deciphering the molecular mechanisms of this disease are therefore needed to develop novel therapeutic approaches.

Here we report the identification of truncated isoforms of the melanoma suppressor TRPM1, generated due to APA followed by premature termination by cleavage and polyadenylation (PCPA). As a second far-reaching consequence of TRPM1 APA we here show reduced miR-211 expression, which is transcribed from a coding sequence located downstream of the identified alternative PAS. This mechanism of APA-dependent regulation of miRNA expression was furthermore confirmed for a second host gene/miRNA pair implicated in melanoma, PUNK1 and miR-107. Both processes, APA and miRNA generation are controlled by U1 expression levels, and decreased U1 abundance in melanoma underscores activation of this newly identified layer of gene expression regulation and its potential pathological impact. Finally, we harnessed the revealed mechanism of dysregulated miRNA expression for a therapeutic approach using antisense morpholino oligonucleotides. Specific targeting of TRPM1 alternative PAS choice not only modulates the expression of TRPM1 protein isoforms, but also inhibits or potentially restores miR-211 expression to endogenous levels by using a single compound.

## Results

### Small RNA sequencing of melanoma cell lines reveals commonly downregulated miRNAs

To identify intronic miRNAs that are commonly deregulated in melanoma, we sequenced small RNAs (18-30 nucleotides) isolated from four selected melanoma cell lines (M14, M19-Mel, SK-MEL-5 und UACC-257) and normal human epidermal melanocytes (NHEM) as control. A miRNA was considered detected if at least 10 read counts per sample were present. We detected 326 miRNAs in the different melanoma cell lines, which were then independently analyzed for differential expression compared to control (Extended Data Table 1). Among these, 28 miRNAs were upregulated and 38 were downregulated with statistically significant differences in all four melanoma cell lines (FDR<0.05, Extended Data Table 2). Out of the 38 downregulated miRNAs, 14 (37%) were located within intragenic regions, with 10 (26%) being located within intronic sequences of protein coding genes (Fig. 1a). 18 (47%) were located within non-coding RNAs, and 4 (11%) within intergenic regions. 2 (5%) had to be considered ambiguous.

**Figure 1:**
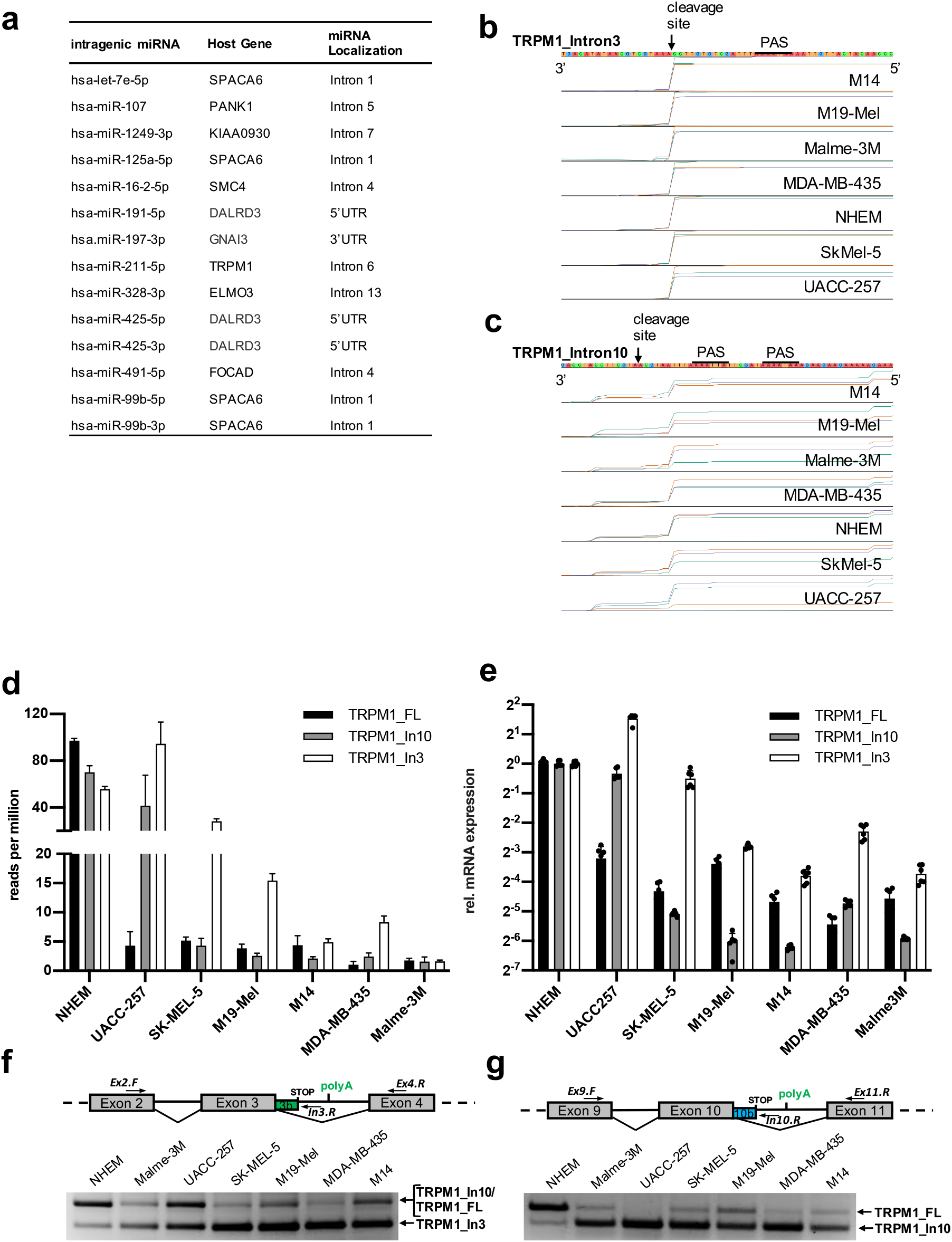
Expression pattern of TRPM1 is decreased and shifted towards novel identified truncated isoforms in melanoma. (a) Small RNA sequencing of melanoma cell lines and NHEM control cells. Differential expression test (log fold change) was performed for all melanoma cell lines in comparison to NHEMs with FDR < 0.05 were considered as differential expressed microRNAs. Common downregulated microRNAs (log fold change < 0) were analyzed for genomic localization. (b-d) QuantSeq 3’mRNA Sequencing of melanoma cell lines and control cell line NHEM. Sequencing data were mapped against human reference genome and visualized in a genome viewer. Section shows human TRPM1 intron 3 (b) and intron 10 (c) sequences. Cleavage site and poly(A) signal are indicated. (d) Reads per million of TRPM1_In3, TRPM1_In10 and TRPM1_FL isoforms. (e) Relative quantification of TRPM1 isoforms were performed by real time PCR with primer pairs specific for each TRPM1 isoform. Expression levels of each isoform in each cell line were normalized to HPRT. Statistical significance was calculated with one-way ANOVA, p<0.05 (*), p<0.01 (**), p<0.001 (***) and p<0.0001 (****). (f) TRPM1 isoform expression patterns were assessed by three-oligo PCR, with a common forward primer (Ex2.F) and two reveres primer, one specific for TRPM1_In3 isoform (In3.R), and one for both, TRPM1_In10 and TRPM1_Fl isoforms (Ex4.R) (g) or with a common forward primer (Ex9.F) and two reveres primer, one specific for TRPM1_In10 isoform (In10.R), and one for TRPM1_FL isoforms (Ex11.R).

To elucidate the molecular mechanism underlying this dysregulation, one of the commonly downregulated miRNAs, miR-211, was chosen as a study target. MiR-211 has previously been shown to act as a tumor suppressor ^21^, and miR-211 expression levels were found to be consistently reduced in melanoma cell lines and melanoma patient samples ^22^. miR-211 is encoded within intron 6 of the calcium permeable ion channel TRPM1. The expression of TRPM1 is inversely correlated with melanoma aggressiveness ^23,24^, and several short protein isoforms that include only the N-terminal part of the protein have been identified in melanocytes and melanoma cells ^25,26^. The exact molecular basis of TRPM1 processing, however, is poorly understood and both alternative splicing or shedding have been proposed as mechanisms ^25,26^.

### Identification of two novel intronic polyadenylation sites in TRPM1 by 3’RNA-sequencing

Given the correlation of truncated TRPM1 peptides with melanoma aggressiveness, we first evaluated whether intronic PAS activation may play a role in the generation of shortened TRPM1 isoforms. To this end, we performed a systematic expressed sequence tag (EST) database screen ^10^, revealing expression of intronic sequences for TRPM1 intron 3 and TRPM1 intron 10 (Extended Data Fig. 1a). To identify polyadenylation and cleavage sites of TRPM1 intron 3 and intron 10, 3’RACE experiments were performed using RNA from UACC-257. Several clones displayed a poly(A) tail following a common cleavage site within intron 3 and intron 10 (Extended Data Fig. 1b). To unbiasedly explore transcription termination, 3’RNA-seq of six melanoma cell lines (M14, M19-Mel, Sk-Mel-5, UACC-257, Malme-3M and MDA-MB-435) and NHEM control cells was performed. This analysis clearly pinpointed 3’-ends with basepair precision at equal positions found in all of the investigated cell lines, identifying an intron 3 isoform (TRPM1_In3, Fig. 1b) and an intron 10 isoform (TRPM1_In10, Fig. 1c), in addition to the full-length TRPM1 isoform (TRPM1_FL, Extended Data Fig. 1c). For both truncated isoforms, putative PAS were located 10-30 nucleotides upstream of the identified cleavage sites. Quantitative analyses of 3’RNA-seq data for TRPM1_In3, TRPM1_In10 and TRPM1_FL revealed low expression of the full-length isoform in all melanoma cell lines, and an increased abundance of intron 10 truncated (UACC-257) and intron 3 truncated (M14, MDA-MB-435, M19-Mel, Sk-Mel-5 and UACC257) TRPM1 transcripts, compared to expression in control cells (Fig. 1d). This expression pattern of low full-length to predominantly truncated TRPM1 isoforms in melanoma cells was validated by quantitative RT-PCR analyses (Fig. 1e). Three-oligo PCRs using a common forward primer and two reverse primers to specifically amplify full-length or truncated isoforms furthermore clearly showed a change in expression patterns of TRPM1_In3 (Fig. 1f) and TRPM1_In10 (Fig. 1g) in all melanoma cell lines compared to NHEM control. Our results demonstrate that TRPM1 pre-mRNA processing is significantly altered in melanoma cell lines, suggesting a shift from full-length to truncated TRPM1 isoforms by activation of the identified PAS in intron 3 and intron 10. Both alternative PAS are located upstream of the first transmembrane domain of TRPM1, and thus are predicted to result in short and soluble TRPM1_In3 (136 aa) and TRPM_10 (409 aa) protein isoforms.

### Alternative PAS choice in TRPM1 is regulated by U1 snRNP levels

U1 snRNP (U1) is an essential component of the spliceosome. However, we and others have shown that U1 also plays an important role in RNA-surveillance by silencing cryptic or alternative PAS within introns ^10,11^. This mechanism termed telescripting protects pre-mRNAs from premature termination by cleavage and polyadenylation (PCPA). To investigate whether U1 participates in the regulation of APA in intron 3 and 10 of TRPM1 (Fig. 2a), we decreased the amount of functional U1 by treating selected melanoma cell lines with an antisense morpholino oligonucleotide that contains a canonical 5’ss sequence (MoU1) and non-targeting control morpholino (MoC). By binding to the region of U1 involved in 5’ss recognition, MoU1 impedes the ability of U1 to recognize 5’ss of pre-mRNAs, thus leading to its functional inactivation. As a control of U1 inhibition, we confirmed reduced inclusion of exon 7 of the SMN2 gene upon MoU1 treatment ^27^ (Extended Data Fig. 2a). qRT-PCR analyses of M19-Mel and UACC-257 cells treated with different MoU1 concentrations demonstrated a dose-dependent shift towards PAS in intron 3 and 10. Although expression of all three isoforms decreased upon U1 depletion due to splicing inhibition, as expected, the full-length isoform decreased to a stronger extent, causing the ratios of intron 3 and intron 10 to full-length transcript levels to increase in both M19-Mel and UACC-257 cells (Fig. 2b, Fig. 2c and Extended Data Fig. 2b). These data demonstrate that U1 exerts regulatory effects on alternative PAS choice in TRPM1 in melanoma cell lines, and that reduced U1 levels expose alternative PAS in intron 3 and intron 10 that activate PCPA and expression of truncated TRPM1 isoforms.

**Figure 2:**
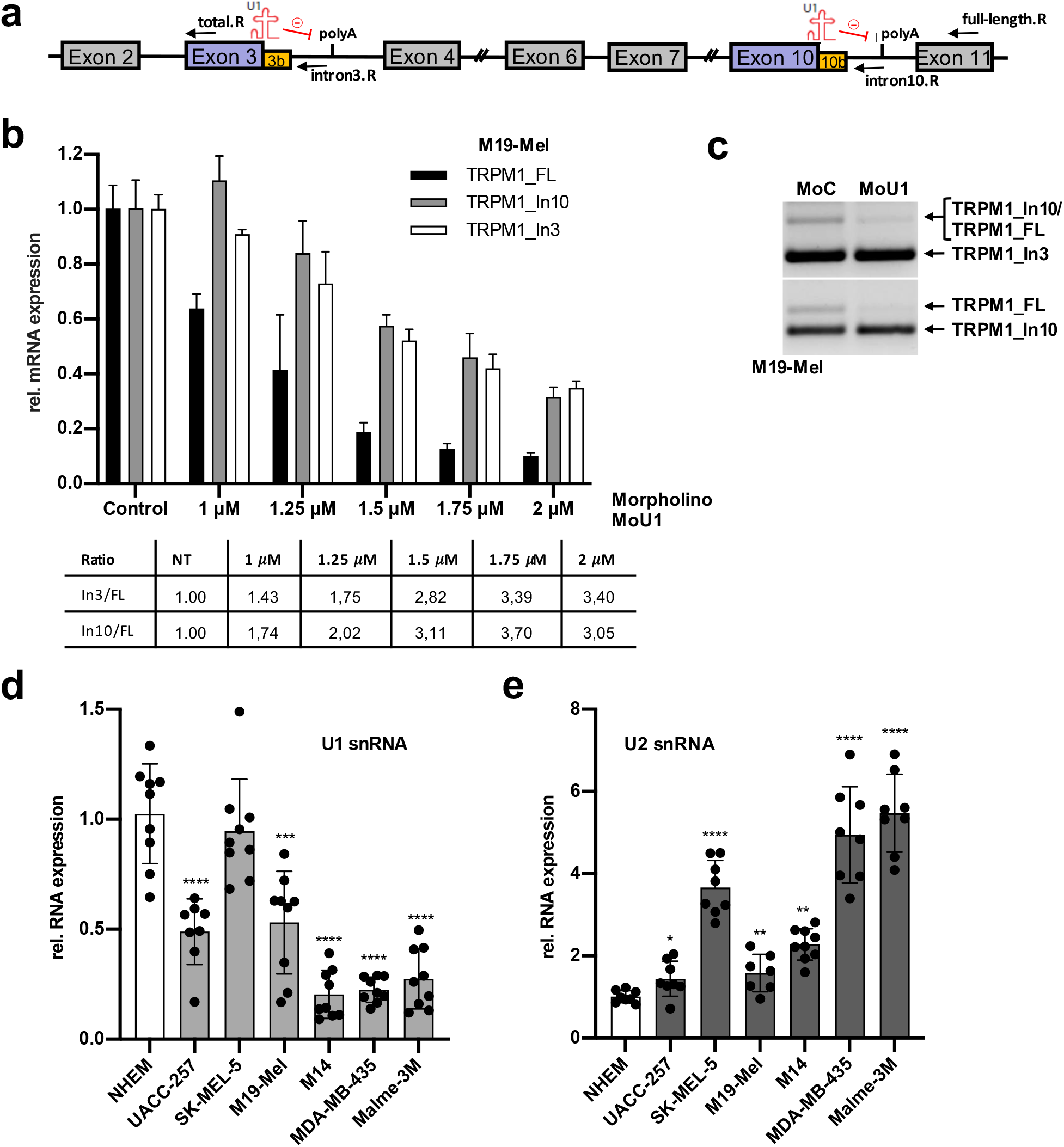
U1 snRNA-dependent alternative intronic polyadenylation of TRPM1 and decreased U1 levels in melanoma cell lines. (a) Scheme representing TRPM1 exon intron structure spanning exon 2 to exon 11. Novel identified alternative poly(A) signals within TRPM1 intron 3 and intron 10 respectively are indicated. Binding of U1 to 5’SS promotes splicing and actively suppresses intronic polyadenylation within TRPM1 intron 3 and intron 10 respectively. (b) Melanoma cell line M19-Mel was treated with indicated concentrations of U1 or control AMO for 10 h. Relative quantification of TRPM1 isoforms were performed by real time PCR with primer pairs specific for each isoform. Expression levels of each isoform were normalized to HPRT and represented relative to control AMO treated cells. Error bars indicate SD. (n=3). Table: Ratios were calculated from fold changes of TRPM1_In3 and TRPM1_FL or TRPM1_In10 and TRPM1_FL. (c) TRPM1 isoform expression patterns were assessed by three-oligo PCR as described in Figure 1 f. (d) Relative quantification of U1 snRNA was performed by real time PCR. Expression levels of U1 snRNA in each cell line were normalized to 18S rRNA and represented relative to NHEM control cells. Error bars indicate SD. (n=9, triplicates from 3 independent cell cultures). Statistical significance was calculated with one-way ANOVA, p<0.05 (*), p<0.01 (**), p<0.001 (***) and p<0.0001 (****). (e) Relative quantification of U2 snRNA were performed by real time PCR. Expression levels of U2 snRNA in each cell line were normalized to 18S rRNA and represented relative to NHEM control cells. Error bars indicate SD. (n=9, triplicates from 3 independent cell cultures). Statistical significance was calculated with one-way ANOVA, p<0.05 (*), p<0.01 (**), p<0.001 (***) and p<0.0001 (****).

Modulation of U1 snRNA levels has previously been shown to regulate cancer cell migration and invasion ^12^. Whether endogenous levels of U1 are altered in melanoma cells, however, has not been investigated. Analysing U1 expression by qRT-PCR in the different melanoma cell lines, we noticed significantly decreased levels in five melanoma cell lines in comparison to NHEMs (Fig. 2d). This indicates changes in the transcriptome since cells of higher organisms produce more U1 than other spliceosomal snRNAs due to a high demand for telescripting ^28^. To rule out a general downregulation of spliceosomal components, we also quantified expression levels of U2 snRNA. Expression of U2 was increased in all melanoma cell lines compared to NHEMs (Fig. 2e), indicating down-regulatory mechanisms specific to U1. Taken together, these results show that U1 blocks activation of intronic poly(A) signals and the generation of truncated TRPM1 isoforms, a mechanism unhinged in melanoma cells, that show decreased U1 levels.

### Intronic polyadenylation of miRNA host genes impacts on miRNA expression

miR-211 is strongly associated with melanoma, and levels of miR-211 are reduced in melanoma patient samples ^21,22^. Given the location of miR-211 within intron 6 of the TRPM1 gene (Fig. 3a), the formation of shortened TRPM1 isoforms in melanoma prompted us to investigate the impact of alternative polyadenylation on miR-211 expression. Our sequencing data of small RNAs had revealed downregulation of miR-211 in melanoma cell lines (Fig. 1a), and miR-211 downregulation could be confirmed in all six melanoma cell lines by qRT-PCR (Fig. 3b and Fig. 3d). Lower miR-211 expression reflects the overall decreased full-length and increased truncated TRPM1 isoform levels, particularly TRPM1_In3, among the 3’RNA-seq reads in respective melanoma cell lines (Fig. 3c).

**Figure 3:**
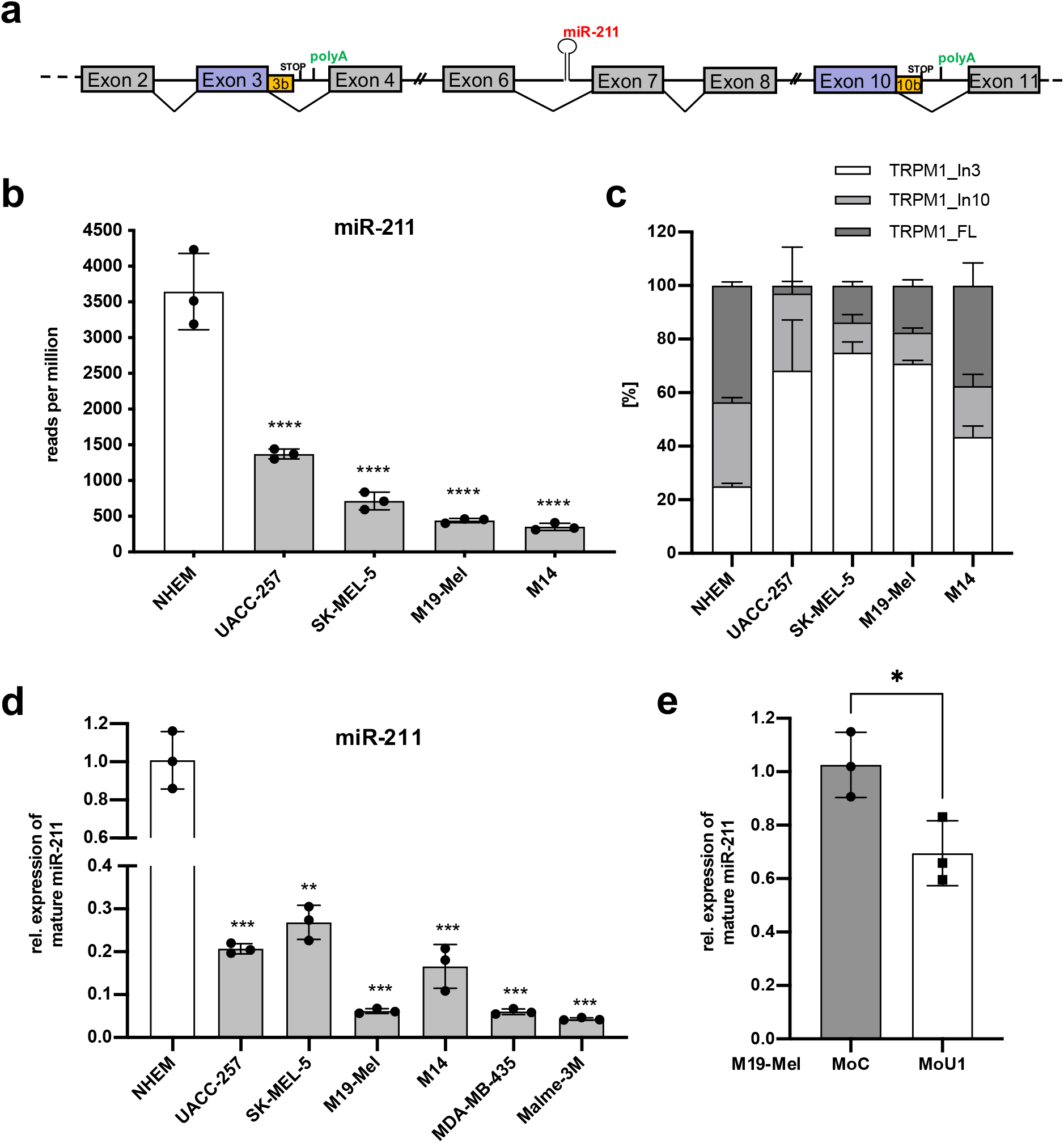
Expression level of intronic miR211 is dependent on host gene TRPM1 isoform expression. (a) Scheme showing TRPM1 exon intron structure spanning exon 2 to exon 11. Novel identified alternative poly(A) signals within TRPM1 intron 3 and intron 10 respectively, as well as intronic microRNA within intron 6 are indicated. (b) Small RNA sequencing of melanoma cell lines and NHEM control cells. Reads per million for hsa-miR-211-5p in each cell line. Statistical significance was calculated with one-way ANOVA, p<0.05 (*), p<0.01 (**), p<0.001 (***) and p<0.0001 (****). (c) Percentage of TRPM1_In3, TRPM1_In10 and TRPM1_FL isoforms in total expression of TRPM1 of the respective melanoma cell line. (d) Relative quantification of mature hsa-miR-211-5p in melanoma cell lines was performed by TaqMan Small RNA Assay. Expression level of hsa-miR-211-5p in each cell line was normalized to RnU6b and represented relative to NHEM control cells. Error bars indicate SD. (n=3). Statistical significance was calculated with one-way ANOVA, p<0.05 (*), p<0.01 (**), p<0.001 (***) and p<0.0001 (****). (e) Melanoma cell line M19-Mel was treated with 1.5 µM of U1 or control AMO for 10 h. Relative quantification of mature hsa-miR-211-5p was performed by TaqMan Small RNA Assay. Expression level of hsa-miR-211-5p was normalized to RnU6b and represented relative to control AMO treated cells. Error bars indicate SD. (n=3). Statistical significant was calculated with Student’s t-test, p<0.05 (*), p<0.01 (**), p<0.001 (***) and p<0.0001 (****).

Importantly, analysis of miR-211 expression in M19-Mel and UACC-257 cell lines after treatment with MoU1 to block U1 binding revealed significantly decreased miR-211 levels (Fig. 3e and Extended Data Fig. 2c), in line with the U1-induced shift towards the truncated TRPM1_In3 isoform terminated upstream of intronic 6, impeding the generation of miR-211. Notably, these results offer a mechanistic explanation for previous unaccounted observations that knockdown of U1 reduces the expression of miR-211 and other intronic but not intergenic miRNAs ^29^.

To show that this model also applies to other host gene/miRNA pairs, we selected miR-107 for further analysis (Fig. 1A). miR-107 was recently described to be downregulated in melanoma and acts as a tumor suppressor ^30^. Its host gene PANK1 is a pantothenate kinase (PANK), which is involved in coenzyme A (CoA) synthesis 31. Our 3’RNA-seq data pinpointed an alternative cleavage site common to all six melanoma cell lines for an intron 1 isoform (PANK1_In1, Extended Data Fig. 3a). A putative PAS is located ∼10 nucleotides upstream of the identified cleavage site and results in a short (121 aa) truncated isoform. Quantitative analyses of 3’RNA-seq data for PANK1_In1 and PANK1_FL revealed lower abundance of the full-length isoform in all investigated melanoma cell lines, and an increased expression of intron 1 truncated PANK1 transcripts, compared to expression in control cells (Extended Data Fig. 3b). miR-107 is located downstream of PANK1_In1 in intron 5, and in accordance with our model, miR-107 is significantly downregulated in all investigated melanoma cell lines as confirmed by small RNA sequencing data (Extended Data Fig. 3c). As for TRPM1, low miR-107 levels compared to control cells reflect a switch in the expression pattern from PANK1 full-length to PANK1_In1 truncated isoform, providing evidence that the interconnection of APA and miRNA expression is a generalizable feature.

### Modulation of alternative polyadenylation and miR-211 expression by antisense morpholino oligonucleotides

Full-length TRPM1 is downregulated in metastatic melanoma and its expression is inversely correlated with metastatic potential and prognosis ^23,24^. Also, miR-211 is widely believed to function as a melanoma tumor suppressor ^21^. Our data strongly indicate a switch towards truncated TRPM1 isoforms and subsequent reduced expression of miR-211 in addition to a transcriptional inhibition of the entire TRPM1 gene in melanoma cells. This opens up the possibility to restore full-length TRPM1 pre-mRNA processing and miR-211 biogenesis using antisense morpholino oligonucleotides. To specifically activate the intronic PAS, we developed a modified splicing re-direction approach using antisense morpholino oligonucleotides (AMOs) ^10^. Typically, AMOs that directly block splice sites cause exon skipping. However, in the presence of putative intronic PAS, AMOs directed against the upstream 5’ss effectively induce PCPA ^10^. Treatment of UACC-257 cells with TRPM1 intron 3 5’ss-blocking morpholino MoMe3 (Fig. 4a, green arrow) indeed led to an upregulation of TRPM1_In3, but decreased levels of TRPM1_FL and TRPM1_In10, indicating activation of the identified alternative intronic PAS (Fig. 4b and Fig. 4c). Of note, analysis of TRPM1 expression by primers amplifying all three isoforms revealed unaltered levels of total transcripts, indicating an exclusive switch from long transcripts to the truncated TRPM1-In3 isoform (Fig. 4b). In an independent approach using HeLa cells transfected with the melanocyte inducing transcription factor (MITF) to induce TRPM1 expression ^26^, MoMe3 treatment reproduced activation of TRPM1_In3 APA (Extended Data Fig. 4a). In accordance with our data revealing an interdependency of APA and miRNA biogenesis, the AMO-induced switch towards TRPM1_In3 was accompanied by a significant decrease in miR-211 levels in UACC-257 cells (Fig. 4d).

**Figure 4:**
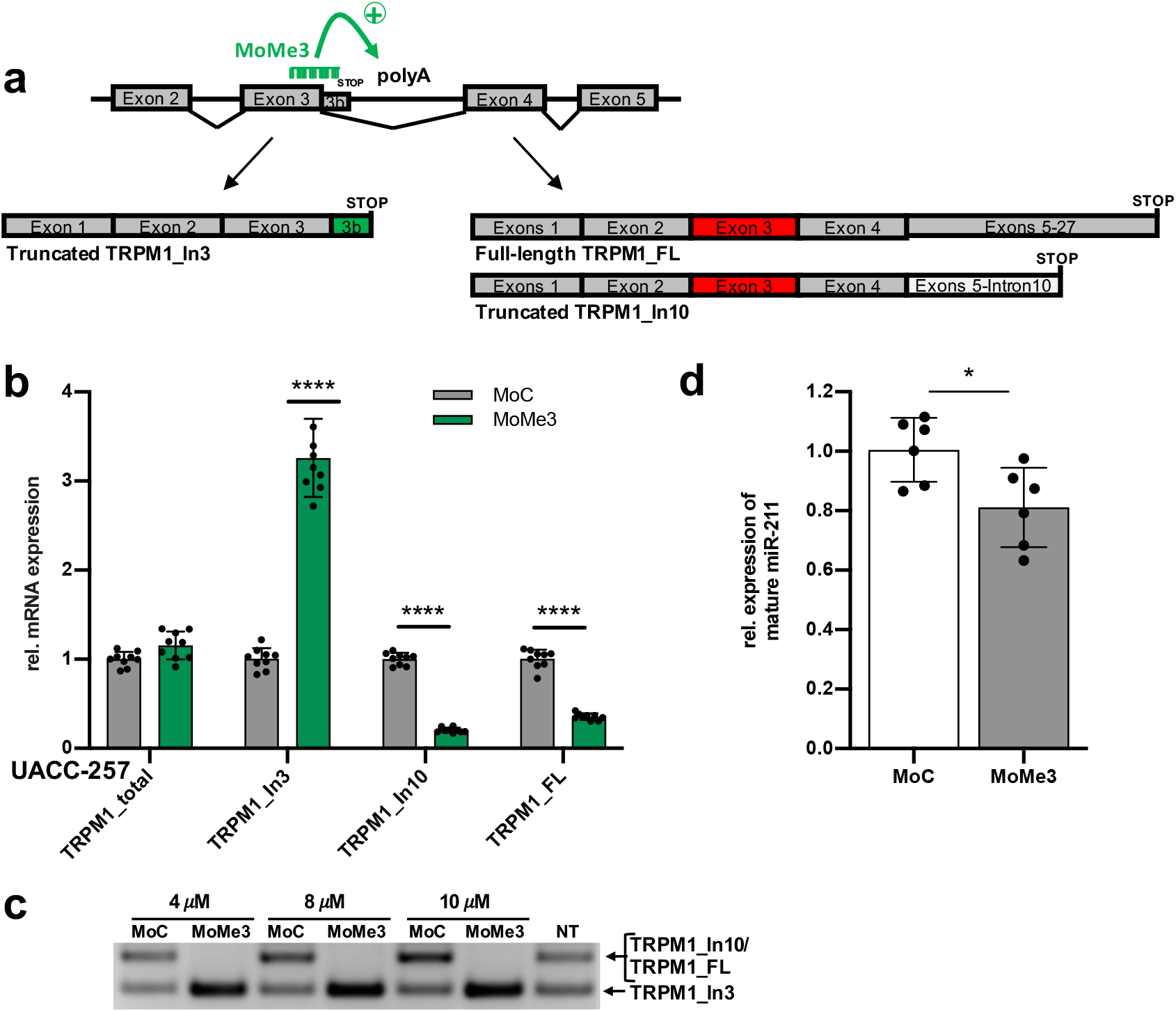
Alternative polyadenylation of TRPM1 can be modulated by antisense oligonucleotides. (a) Scheme representing TRPM1 gene exon intron structure spanning exon 2 to 5. Poly(A) signal in TRPM1 intron 3 is indicated. MoMe3, targeted to TRPM1 exon 3 intron 3 junction (indicated in green), blocks the 5’SS and leads to the activation of intron 3 IPA and the generation of truncated TRPM1_In3 isoform. MoMepA, targeted to novel identified intron3 poly(A) signal (indicated in red), blocks intron 3 IPA and leads to the generation of TRPM1_In10 and TRPM1_FL isoforms. (b-d) Melanoma cell line UACC-257 was treated with 10 µM MoMe3 for 72 h. (b) Relative quantification of TRPM1 isoforms were performed by real time PCR with primer pairs specific for each isoform. Expression levels of each isoform were normalized to HPRT and represented relative to control AMO treated cells. Error bars indicate SD. (n=9) (c) Relative quantification of mature hsa-miR-211-5p was performed by TaqMan Small RNA Assay Expression level of hsa-miR-211-5p was normalized to RnU6b and represented relative to control AMO treated cells. Error bars indicate SD. (n=6). (d) TRPM1 isoform expression patterns were assessed by three-oligo PCR as described in Fig. 1f.

### Expression pattern in primary melanoma tumor samples shows a shift towards truncated TRPM1_3 isoform

Full-length TRPM1 has been shown to be downregulated in metastatic melanoma ^23,24^, and several short TRPM1 protein isoforms have been identified in melanocytes and melanoma cells ^25,26^. We evaluated TRPMI transcript expression in primary melanoma patient samples by three-oligo PCR (Fig. 5a). Although expression levels were heterogeneous across patients, an overall increase in TRPM1_In3 relative to full-length/TRPM1_In10 isoforms (and compared to NHEM controls) was observed (**Figure 5B**). This suggests modulation of TRPM1 pre-mRNA processing and a shift in the expression towards the truncated intron 3 TRPM1 isoform also in melanoma patient samples.

**Figure 5:**
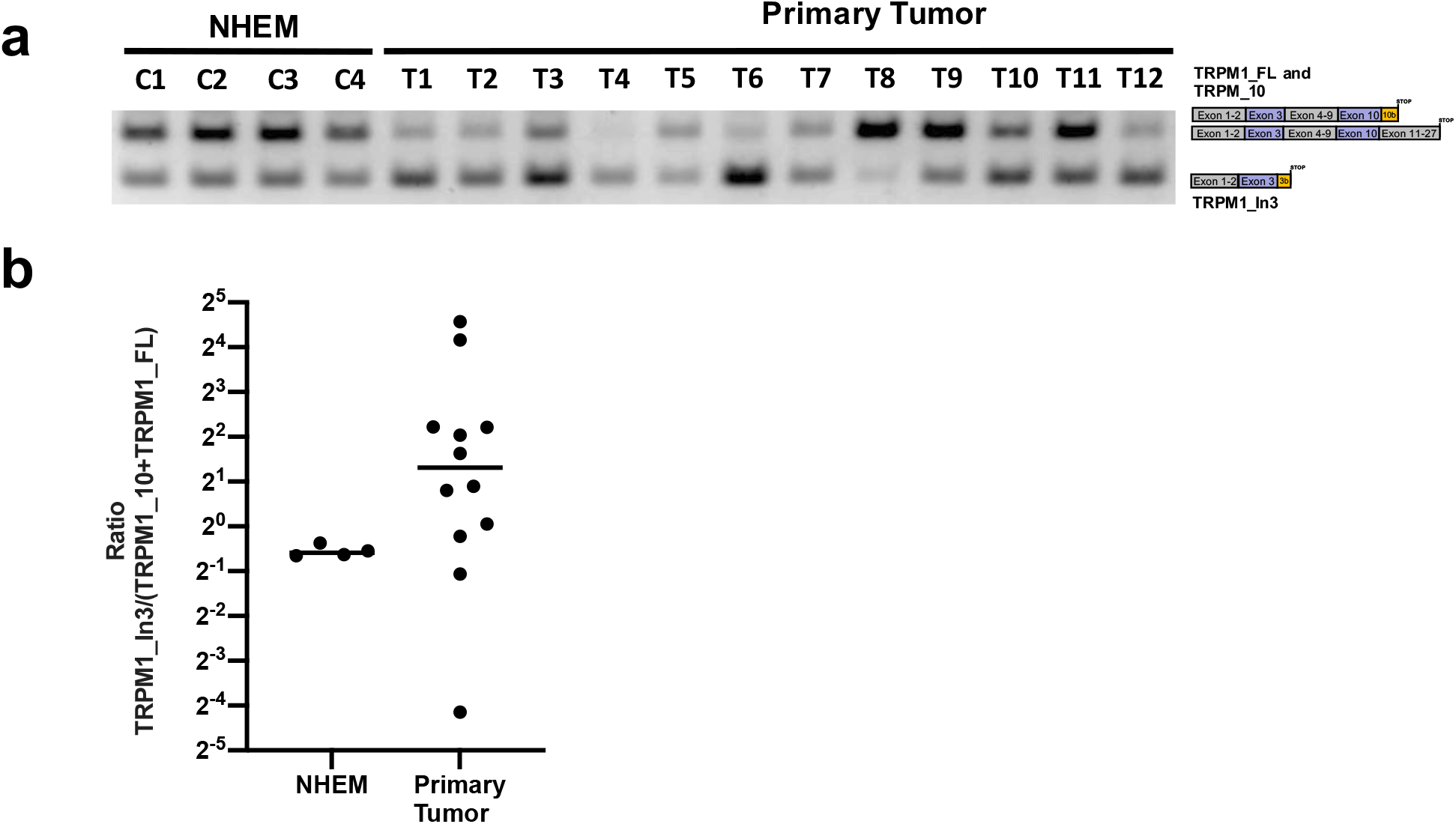
Primary melanoma tumor samples show an expression pattern shift towards truncated TRPM1_In3 isoform. (a) Malignant melanoma patient samples obtained from primary tumor (T1-T12) were analyzed by three oligo PCR as described in Figure 1F. (n=12) (b) Quantification of agarose gel bands shown in (a) by ImageQuant.

Overall, we here show that perturbations in U1 levels in melanoma cell lines resulted in PCPA and an increased expression of a truncated intron 3 TRPM1 isoform. Termination of the transcript upstream of the miRNA coding sequence in addition led to a decrease in miR-211 expression. Our results thus reveal an unexpected additional layer of U1 function exceeding its well-known role in splicing, which controls the expression of truncated protein isoforms as well as miRNA transcript levels. Both of these mechanisms may contribute to the U1 tumor suppressor-like activity of U1 ^12^. We anticipate that targeting these U1-dependent functions could open up new strategies for novel antisense based therapeutic approaches in cancer and other diseases to modulate APA and miRNA with a single, gene specific compound.

## Discussion

APA has emerged as an important player influencing the diversity of gene expression and has been implicated in various human diseases. In this study, we expand our understanding of pathological outcomes of APA beyond the generation of short protein isoforms and the deletion of miRNA binding sites within the 3’UTR. Instigation of APA can have effects far beyond the gene in which it occurs: By blocking the biogenesis of intronic miRNAs, a single APA event can affect the multitude of target genes of these miRNA, having potentially genome wide consequences.

Both, APA and miRNA expression are frequently deregulated in human diseases such as cancer, CNS disorders, inflammation, cardiovascular diseases and metabolic disorders ^4-7^. The fact that more than half of all miRNAs are encoded within intronic regions of protein coding genes suggests that these processes are coupled. So far, however, events leading to changes in pre-mRNA-processing or to miRNA-expression have mostly been studied separately. Integral approaches focusing on both pre-mRNA and miRNA regulation at the same time are scarce. For example, despite recognizing the emerging role of APA in gene expression regulation, many studies focused on the loss of miRNA binding sites within the 3’UTR ^32,33^ rather than consequences of intronic APA beyond shortened transcripts ^34^.

In the context of cancer, it has been demonstrated that highly proliferative and poorly differentiated cells often produce shorter and proximally polyadenylated isoforms ^4,35^, and that the resulting truncation of 3’UTRs reduces the number of miRNA-binding sites.^9^ Thus, increased APA interferes with the tightly controlled management of protein expression by RNA-interference observed in normal cells. However, this mechanism only affects the regulation of the protein affected by APA. Beyond this, we here show that activation of intronic polyadenylation due to newly identified poly(A) sites not only leads to a shift from full-length to truncated transcripts but also to a concomitant downregulation of their hosted miRNAs. By contrast, this mechanism affects the expression of other genes, namely those with binding sites for the eliminated miRNAs, rather than of the truncated host gene, and may contribute to the dysregulation of miRNAs in cancer beyond the previously described processes. The revealed mechanism might also explain why miRNAs tend to be downregulated in transformed cells ^36,37^, despite the fact that expression of protein-coding genes tends to be upregulated in highly proliferative cells. Aside from cancer, it has recently been shown that a genome-wide shift to short mRNA transcripts is the predominant response to heat shock and induced by APA ^8^. A possible so far unresolved biological function of this stress-induced shift could be the global and rapid shut down of miRNA generation to protect transcripts from degradation. We here were able to demonstrate downregulation of intronic miRNAs after activation of APA of two genes involved in the pathology of melanoma, TRPM1 and PANK1, and their hosted miRNAs miR-211 and miR-107 respectively, as examples. The explicit connection between truncated pre-mRNA by APA and loss of miRNA expression point towards a general mechanism.

In line with U1’s function in telescripting ^10,11^, we were able to induce a shift from full-length to truncated isoforms by inhibiting U1 base-pairing with morpholino antisense oligonucleotides. However, previous studies show numerous and diverse transcriptomic changes when U1 levels are experimentally modulated, including upregulation of oncogenes and downregulation of tumor suppressor genes, leading to an invasive and migratory phenotype of cancer cells ^12^. Also, the UV-induced DNA damage response correlates with decreases levels of U1 and APA ^38^. Whether inhibition of U1 leads to changes in miRNA expression levels and may serve as an explanation for the numerous transcriptomic changes under these circumstances has not been investigated. Our results show that inhibition of U1 not only leads to intronic APA in TRPM1 but also results in lower levels of miR-211, linking U1 to miRNA expression. Additionally, significantly lower levels of U1 in melanoma cells suggest a fundamental role of U1 in malignancies: its downregulation in cancer may spur an escape from cellular expression control by reducing potential miRNA binding sites within the 3’ UTR in the APA-dysregulated gene, but also decrease miRNA expression from intronic sites affecting genome wide protein expression. An interesting question remains, i.e. if the observed U1 downregulation in cancer is a result of the malignant transformation, or affects carcinogenesis itself, as suggested by the immediate decrease in U1-levels upon UV-damage ^38^.

Another important question is if the identified novel mechanism behind miRNA-regulation can be exploited therapeutically. As miRNAs are dysregulated in various diseases, considerable efforts currently focus on the development of therapeutics to modulate miRNA activity ^18,39^. For example, miRNA mimics aim to increase levels of a miRNA to endogenous levels. Potential off-target effects are due to uptake by non-target tissues and the introduction of supraphysiologial levels in target cells ^40^. By blocking the 5’ss of TRPM1 using an antisense morpholino oligonucleotide, we were able to inhibit miR-211 biogenesis by shifting the APA pattern from full-length to the truncated isoform. Modulating miRNA expression by targeting APA of its host gene ensures therapeutic miRNA dosing close to its endogenous levels. By this new strategy, microprocessing of a tumor suppressive miRNA can be restored by blocking alternative poly(A) signals or *cis*-regulatory elements of its host gene, with concomitant modulation of protein isoform expression patterns using a single antisense compound.

In this way, our results not only add to the already complex regulation of pre-mRNA processing and miRNA biogenesis. They also offer the possibility of a novel therapeutic approach to restore normal miRNA activity in a wide spectrum of diseases.

## Methods

### Cell culture

Melanoma cells (M14, MDA-MB-435, M19-Mel, SK-Mel-5, UACC-257, Malme-3M obtained from the Department of Dermatology, University Hospital Wuerzburg) were maintained in RPMI1640 medium supplemented with 10% FBS (Sigma Aldrich) and 10 units/ml penicillin and 10 μg/ml streptomycin (Thermo Fisher Scientific) at 37 °C and 5% CO_2_. Normal human epidermal melanocytes (NHEM, obtained from PromoCell) were maintained in complete Melanocyte Growth Medium M2 (PromoCell) at 37 °C and 5% CO_2_.

### Morpholino Treatment

U1 antisense and control morpholino oligonucleotide sequences were GGTATCTCCCCTGCCAGGTAAGTAT and CCTCTTACCTCAGTTACAATTTATA respectively^11^. Oligonucleotides are coupled to a delivery moiety (vivo-morpholino, Gene Tools) and were directly added to 4 × 10^4^ cells according to the manufacturer’s instruction to achieve the desired final doses indicated in the text and figures. MoMepA, MoMe3 and control morpholino oligonucleotide sequences were TTTATTTTAACAATGATGTTGGGCC, ATGTCTGAATGCCTTTCTCACCATG and CCTCTTACCTCAGTTACAATTTATA respectively. Oligonucleotides were transfected into 4 × 10^4^ cells using Endoporter (Gene Tools) according to the manufacturer’s instruction to achieve the desired final doses indicated in the text and figures.

### Melanoma samples

Snap frozen melanoma samples were obtained after written informed consent from patients undergoing resection of primary tumors. Prospective collection was approved by local ethics committee (study reference: 241/2014). Samples were collected between September 2015 and December 2016 at the Department of Dermatology, University Hospital Wuerzburg and stored at -80°C until usage.

### RNA Isolation, RT-PCR and quantitative PCR

Total RNA of cells was isolated with Nucleo Spin miRNA Kit (Macherey-Nagel), total RNA of melanoma samples was isolated with RNeasy Kit (Qiagen) according to the manufacturer’s instruction. 500 ng total RNA were reverse transcribed with cDNA first strand synthesis kit (Thermo Fisher Scientific) either with Random or oligo(dT) primer.

Three-oligo PCR was performed with 2xPCR MasterMix (Invitrogen) with a common forward primer (Ex2.F) and two reverse primers, one specific for TRPM1_In3 (In3.R), and one for both, TRPM1_In10 and TRPM1_FL (Ex4.R) or with a common forward primer (Ex9.F) and two reverse primers, one specific for TRPM1_In10 (In10.R) and one for TRPM1_FL (Ex11.R).). All primer sequences are summarized in Extended Data Table 3.

Quantitative PCR reactions for TRPM1 isoforms and U1 were prepared with Power up SYBR Green MasterMix (Invitrogen) and carried out on Quant Studio 6 Flex System (Applied Biosystems). Relative gene expression was calculated by the 2−ΔΔCT method. HPRT or 18S rRNA were used as the internal control for normalization. Reverse Transcription and quantitative PCR of mature miR-211 from total RNA was performed with TaqMan small RNA Assay (Applied Biosystems) according to the manufacturer’s instruction and carried out on Quant Studio 6 Flex System (Applied Biosystems). Relative microRNA expression was calculated by the 2−ΔΔCT method and RnU6b was used as the internal control for normalization.

### Preparation and sequencing of QuantSeq 3’ mRNA-Seq REV libraries

Preparation of QuantSeq 3’ mRNA-Seq REV libraries and sequencing was performed by the Core Unit SysMed University Wuerzburg, Briefly, sequencing libraries were prepared from total RNA using the QuantSeq 3′ mRNA-Seq Library Prep Kit REV for Illumina (Lexogen) according to manufacturer’s instruction. Paired end sequencing

(2x 75bp) of libraries was performed on an Illumina Nextseq 500. Reads were mapped against the human reference genome (Ensemblversion 90) using STAR 2.5.3a ^41^

### Preparation and sequencing of microRNA libraries

Preparation and sequencing of microRNA libraries was performed by vertis biotechnology AG. Briefly, oligonucleotide adapters were ligated to the 5’ and 3’ ends of the RNA samples. First-strand cDNA synthesis of total RNA was performed using M-MLV reverse transcriptase and the 3’ adapter as primer. The resulting cDNAs were amplified with PCR using a high fidelity DNA polymerase. miRNA cDNA was pooled and fractionated in the size range of 150 - 170 bp using a polyacrylamide (PAA) gel. cDNA pools were sequenced on an Illumina NextSeq 500 system using 75 bp read length. Trimmed reads were mapped against miR database release 22 (human).

### Identification of intronic microRNA and putative intronic APA sites

The miRNA counts table was used as input for EdgeR ^42^ after filtering low counts miRNAs (at least 10 counts in each sample for a miRNA to be retained). Raw counts were normalized with the TMM function of EdgeR. P values for differential expression were computed for each melanoma cell line compared to NHEM based on a generalized linear model with NHEMs as baseline (Quasi-Likelihood F-test). P values were corrected for multiple testing by the Benjamini-Hochberg method. Differentially expressed miRNAs were sorted by false discovery rate (FDR) and subsequently a 0.05 FDR threshold was used to filter each set. Only miRNAs that were present in all filtered sets were kept for the following steps.

To define intronic miRNAs a GTF file was downloaded from Ensembl (version 102) and the latest miRNA human GFF3 annotation was downloaded from miRbase (version 22.1). Using Bedtools^43^ a BED format file was produced from each annotation file and the Intersect command was used to find the genomic overlap. miRNAs that were included in this overlap were considered intronic.

## Data availability

The data for this study have been deposited in the European Nucleotide Archive (ENA) at EMBL-EBI under accession number PRJEB51529 (https://www.ebi.ac.uk/ena/browser/view/PRJEB51529). All RNA-seq miRNA sequencing data in this publication have been deposited in the Gene Expression Omnibus with the accession number GSE 199181.

## Code availability

Code for the analyses described in this study is available from the corresponding author upon request.

## Acknowledgments

This project was supported by the Interdisciplinary Center for Clinical Research (Interdisziplinäres Zentrum für Klinische Forschung (IZKF)), University Hospital Würzburg (E-352 to S.V. and A.Z. and A-384 to A.Z.), the Deutsche Forschungsgemeinschaft (DFG; German Research Foundation, 374031971 - TRR 240, 324392634 - TR221, ZE827/13-1, 14-1, project numbers 396923792 and 432915089 to A.Z.) and the Fritz Thyssen Stiftung (Az. 10.17.1.025MN to S.V.).

## Author Contributions

Conceptualization, S.V. and A.Z.; Investigation, G.B., Y.K., A.R., M.B., M.E., E.H. and E.B.; Formal Analysis, R.B., L.D., F.E.; Writing – Original Draft, S.V. and A.Z. with input from all authors; Funding Acquisition, S.V. and A.Z.; Resources, E.H., R.H., B.S., F.E., U.F.; Supervision, S.V. and A.Z.;

## Declaration of Interest

The authors declare no competing interests.

## Notes

### Competing Interest Statement

The authors have declared no competing interest.

