## Supplemental Data for "U1 snRNP suppresses microRNA biogenesis by alternative intronic polyadenylation in melanoma"

**Extended Data Figures 1-4 and Table S3** **Page**

| | | |
|---|---|---|
| • Extended Data Fig. 1 | Identification and validation of novel truncated human TRPM1 isoforms | S2 |
| • Extended Data Fig. 2 | Additional confirmation of the impact of U1 snRNA on alternative intronic polyadenylation | S3 |
| • Extended Data Fig. 3 | Expression level of intronic miR107 is decreased in melanoma cell lines due to a shift towards truncated isoforms of host gene PANK1 | S4 |
| • Extended Data Fig. 4 | TRPM1 expression can be induced in HeLa cells by MITF overexpression and be modulated by AMOs | S5 |
| • Extended Data Table 3 | Primer Sequences | S6 |

#### Extended Data Fig. 1

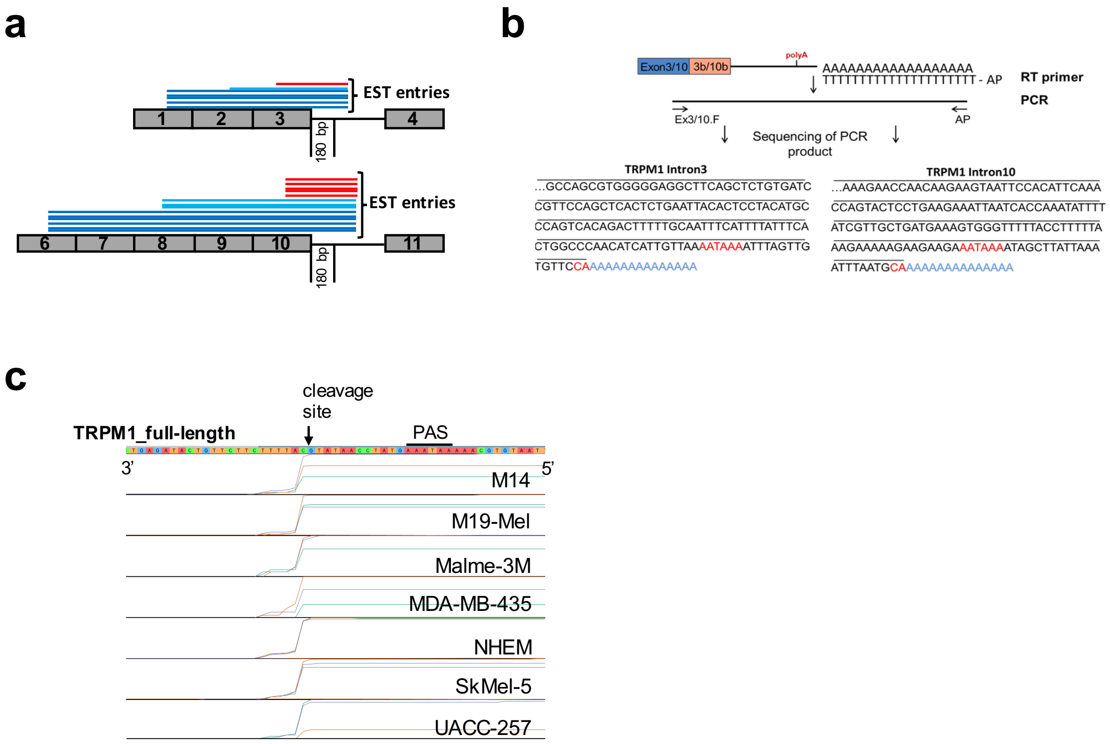


**Extended Data Fig. 1: Identification and validation of novel truncated human TRPM1 isoforms**

**(**a) The first 180 bps of TRPM1 introns 1 to 13 were used to query EST databases with BLASTN. Positive hits (score > 80) were further analyzed. (B) Retrieved GenBank EST hit entries were aligned to the TRPM1 genomic sequences. Hits that ended within the first upstream exon were considered “weak” (in red) since neither upstream processing nor unprocessed RNA can be excluded. Hits that aligned to the exon-intron boundary and spanned one or more upstream exon-exon junctions, indicating evidence of processed RNAs were considered “strong” positive hits (in blue). (b) Schematic showing TRPM1 exon 3/10 and intron 3/10 with putative intronic PAS. For 3’RACE, reverse transcription reaction was performed with an oligo(dT)-anchor primer, followed by PCR with a reverse primer corresponding to the adaptor sequence and a forward primer specific to TRPM1 exon 3 or exon 10 respectively. Amplified PCR products were directly sequenced after gel extraction. Partial cDNA sequences of the PCR fragments for TRPM1 intron 3 or intron 10 respectively. PAS and cleavage sites are indicated by red letters, poly(A) tail in blue. (c) Relative quantification of TRPM1 isoforms were performed by real time PCR with primer pairs specific for each isoform. Expression levels of each isoform in each cell line were normalized to NHEM control cells. Error bars indicate SD. (n=9).

#### Extended Data Fig. 2

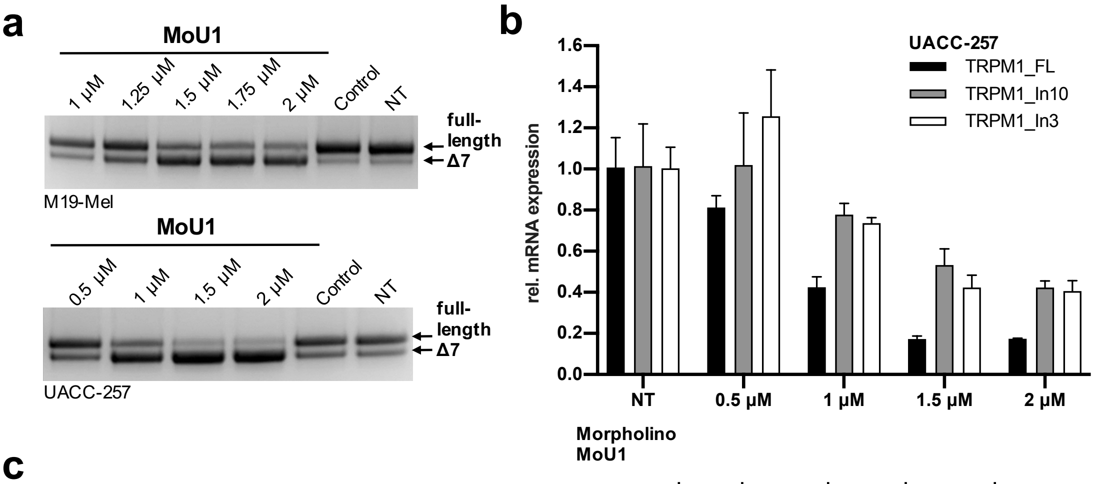


| Ratio | NT | 0.5 μM | 1 μM | 1.5 μM | 2 μM |
|---|---|---|---|---|---|
| In3/FL | 1.00 | 1,56 | 1,75 | 2,46 | 2,16 |
| In10/FL | 1,02 | 1,27 | 1,84 | 3,10 | 2,33 |

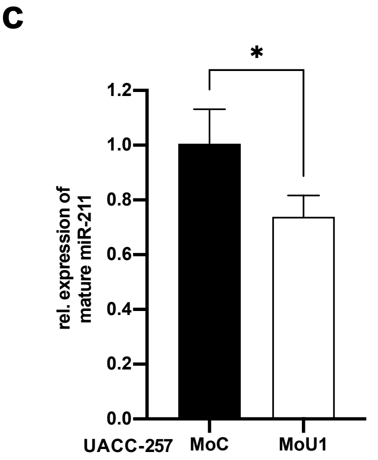


**Extended Data Fig. 2: Additional confirmation of the impact of U1 snRNA on alternative intronic polyadenylation**

(a) Functional U1 snRNA depletion by increasing U1 AMO concentrations was analyzed by PCR for reduced inclusion of exon 7 of the SMN2 gene in melanoma cell lines UACC257 and M19-Mel. (b) Melanoma cell line UACC257 was treated with indicated concentrations of U1 or control AMO for 10 h. Relative quantification of TRPM1 isoforms were performed by real time PCR with primer pairs specific for each isoform. Expression levels of each isoform were normalized to HPRT and represented relative to control AMO treated cells. Error bars indicate SD. (n=3). Table: Ratios were calculated from fold changes of TRPM1_In3 and TRPM1_FL or TRPM1_In10 and TRPM1_FL. (d) Relative quantification of mature hsa-miR-211-5p after treatment with U1 or control AMO in melanoma cell line UACC257 (1.5 μM, 10 h) was performed by TaqMan Small RNA Assay. Expression level of hsa-miR-211-5p was normalized to RnU6b and represented relative to control AMO treated cells. Error bars indicate SD. (n=3). Statistical significance was calculated with Student's t-test, p<0.05 (*), p<0.01 (**), p<0.001 (***) and p<0.0001 (****).

#### Extended Data Fig. 3

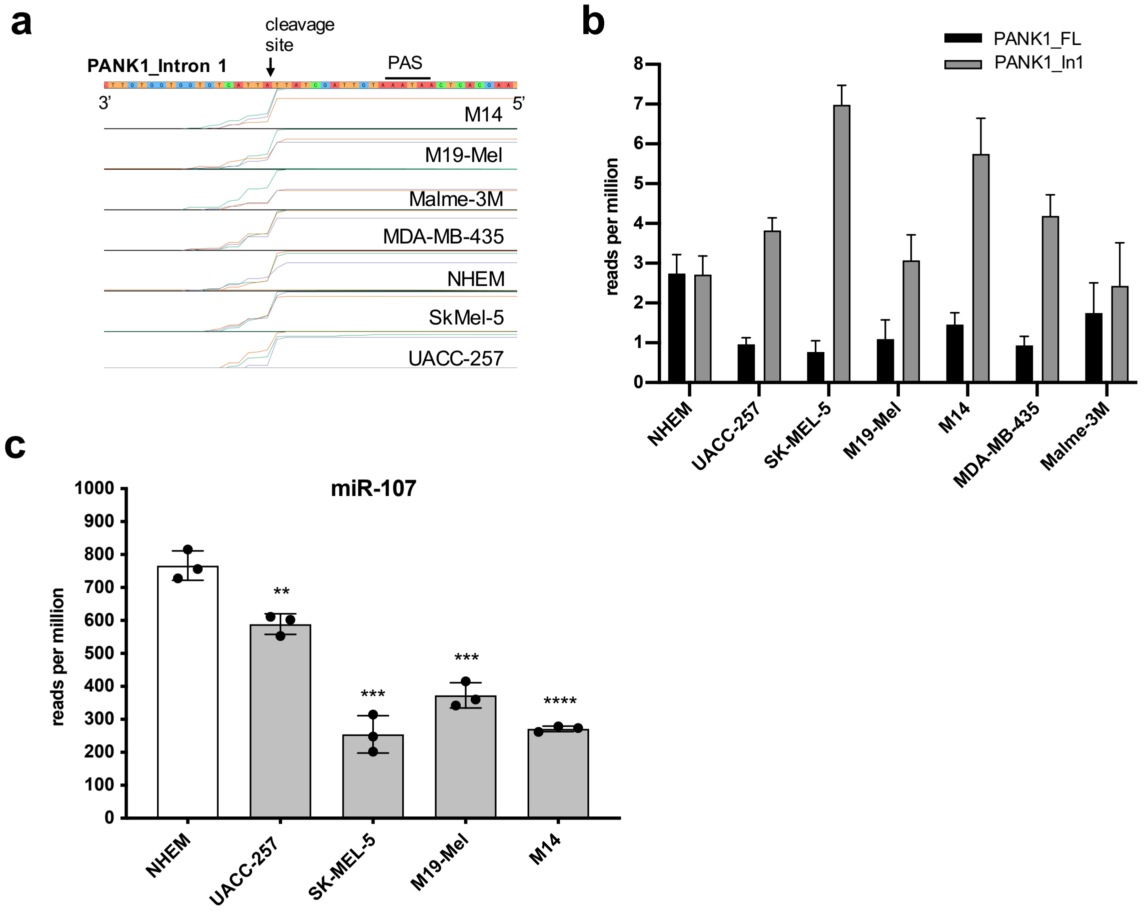


**Extended Data Fig.3: Expression level of intronic miR107 is decreased in melanoma cell lines due to a shift towards truncated isoforms of host gene PANK1.**

(a) QuantSeq 3'mRNA Sequencing of melanoma cell lines and control cell line NHEM. Sequencing data were mapped against human reference genome and visualized in a genome viewer. Section shows human PANK1 intron 1 sequence. Cleavage site and poly(A) signal are indicated. (b) Reads per million of PANK1_In1 and PANK1_FL isoforms. (c) Small RNA sequencing of melanoma cell lines and NHEM control cells. Reads per million for hsa-miR-107 in each cell line. Statistical significance was calculated with one-way ANOVA, $p<0.05$ (*), $p<0.01$ (**), $p<0.001$ (***) and $p<0.0001$ (****).

### Extended Data Fig. 4

a

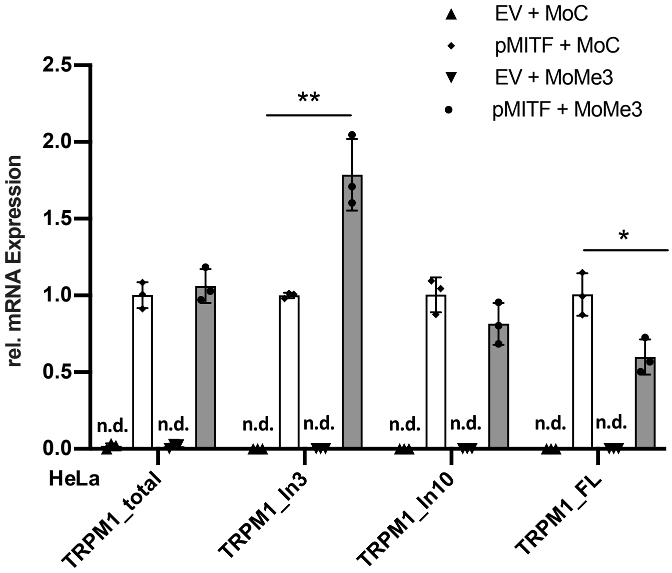


**Extended Data Fig. 4: TRPM1 expression can be induced in HeLa cells by MITF overexpression and be modulated by AMOs.**

(a) HeLa cells were treated with 10 µM of MoMe3 or Ctrl AMO for 4 h. Afterwards cells were transfected with pMITF or empty vector (EV) for 48 h. Relative quantification of TRPM1 isoforms were performed by real time PCR with primer pairs specific for each isoform. Expression levels of each isoform were normalized to HPRT and represented relative to control AMO treated and pMITF transfected cells. Error bars indicate SD. (n=3).

**Extended Data Table 1:** 326 detectable miRNAs' raw counts. All the miRNAs present with in at least 10 counts in all cell lines were used for the differential expression analysis, after normalization with the TMM method of EdgeR.

**Extended Data Table 2:** 66 differentially expressed miRNAs in melanoma cell lines with consistent effect direction. For each comparison to the NHEM cell line, miRNAs were sorted by FDR were selected. After filtering for FDR < 0.05 the miRNAs that were found to be consistently down-regulated (38, red text) or up-regulated (28, in blue) were retained. 24 miRNAs were annotated as intronic.

**Extended Data Table 3:** Primer Sequences

| Target | Sequence |
|---|---|
| TRPM1 Intron3 | Forward: GTAATTCCTAGCATGAAAGACTCTAACAG<br>Reverse: CTGTGATCCTAGTGGCACCA |
| TRPM1 Intron10 | Forward: TTCAGAATGGGTTCTGAGGGC<br>Reverse: TTCCATGTCACAAGGGAGAACAT |
| TRPM1 FullLength | Forward: TCTCGTGGCCATTTCCACAT<br>Reverse: TTTCGGGCCAGTTTCCAAGA |
| TRPM1 Total | Forward: GTAATTCCTAGCATGAAAGACTCTAACAG<br>Reverse: TTGGAATATCCGCCACCCTG |
| HPRT | Forward: TGACCTTGATTTATTTTGCATACC<br>Reverse: CGAGCAAGACGTTCAGTCCT |
| 18S rRNA | Forward: GGCCCTGTAATTGGAATGAGTC<br>Reverse: CCAAGATCCAACTACGAGCTT |
| U1 snRNA | Forward: TGATCACGAAGGTGGTTTTCC<br>Reverse: CATTGCACTCCGGATGTGC |
| U2 snRNA | Forward: CTCGGCCTTTTGGCTAAGAT<br>Reverse: TGTCCTCGGATAGAGGACGTA |
| U6 snRNA | Forward: GCTTCGGCAGCACATATACTA<br>Reverse: CGCTTCACGAATTTGCGTGTC |
